# Perfect mimicry between *Heliconius* butterflies is constrained by genetics and development

**DOI:** 10.1101/2020.01.10.902494

**Authors:** Steven M. Van Belleghem, Paola A. Alicea Roman, Heriberto Carbia Gutierrez, Brian A. Counterman, Riccardo Papa

## Abstract

Müllerian mimicry strongly exemplifies the power of natural selection. However, the exact measure of such adaptive phenotypic convergence and the possible causes of its imperfection often remain unidentified. The butterfly species *Heliconius erato* and *Heliconius melpomene* have a large diversity of co-mimicking geographic races with remarkable resemblance in melanic patterning across the mid-forewing that has been linked to expression patterns of the gene *WntA*. Recent CRISPR/Cas9 experiments have informed us on the exact areas of the wings in which *WntA* affects color pattern formation in both *H. erato* and *H. melpomene*, thus providing a unique comparative dataset to explore the functioning of a gene and its potential effect on phenotypic evolution. We therefore quantified wing color pattern differences in the mid-forewing region of 14 co-mimetic races of *H. erato* and *H. melpomene* and measured the extent to which mimicking races are not perfectly identical. While the relative size of the mid-forewing pattern is generally nearly identical, our results highlight the areas of the wing that prevent these species from achieving perfect mimicry and demonstrate that this mismatch can be largely explained by constraints imposed by divergence in the gene regulatory network that define wing color patterning. Divergence in the developmental architecture of a trait can thus constrain morphological evolution even between relatively closely related species.

## 1. Introduction

Adaptation is the product of natural selection as well as the ability of a population to generate adaptive genetic diversity for natural selection to act on [1]. In this regard, irreversible steps at key developmental stages can limit or bias the production of variant phenotypes, posing so-called developmental constraints on the evolution of phenotypes [2]. Moreover, when populations evolve independently, divergence at these steps can also lead evolution along an irreversible trajectory [3]. Understanding the relative contribution of genetics and development to adaptation would therefore allow us to better understand the directionality and predictability of evolution [4]. However, due to the difficulties of studying the role of genetics and development in generating phenotypic variation, to date, few evolutionary study systems have been able to assess the interplay of natural selection with genetic and developmental mechanisms.

Cases of convergent evolution in distinct lineages provide powerful opportunities to investigate the selective, genetic and developmental routes to adaptation [1]. For example, convergent evolution between co-mimetic butterfly species provides a comparative framework to investigate the genes that are co-opted in the evolution of a trait, including their regulation and interactions with other factors [5–7]. Particularly, in the case of Müllerian mimicry, in which both partners have evolved an honest aposematic warning signal, natural selection is expected to strongly favor convergence to the same color pattern [8,9]. However, the genetic changes and molecular mechanisms driving such phenotypic convergence are less obvious. Therefore, quantifying the extent of wing color pattern resemblance between Müllerian mimics could allow us to investigate the interplay of selective, genetic and developmental mechanisms underlying convergent evolution. Such information is necessary in order to understand the potential existence of developmental constraints that might limit the potential of selection and the feasibility of repeated outcomes in evolution.

Mimicry between the two butterfly species *Heliconius erato* and *Heliconius melpomene* has long been a key illustration of the perfecting power of natural selection [10,11]. In these Müllerian mimics, positive frequency dependent selection imposed by birds has favored the evolution of over 25 geographically distinct mimetic populations of unpalatable butterflies. Although there is no evidence for gene flow between *H. erato* and *H. melpomene*, which split around 12-14 Mya [12], their resemblance in wing colour patterns is remarkable. For example, their mid-forewing colour pattern shape exhibits incredible diversity within each species yet qualitatively identical morphologies between each co-mimetic population. Genetic research has demonstrated that most of the complexity of color pattern variation in this diverse genus is controlled by only a handful of loci acting broadly across the fore- and hindwings [6,13–16]. These genes have been shown to be repeatedly involved in both the evolution of divergent and convergent phenotypes in *Heliconius*, as well as other butterfly and moth species [7,14,17]. What has been suggested to define the variability in wing color patterns in *Heliconius*, despite the few genes involved, is a complex array of *cis*-regulatory regions that control expression during their wing development [16,18–20].

Recently, a series of functional experiments have knocked out *WntA* in *H. erato* and *H. melpomene* [5,21], a gene that codes for a ligand involved in the gene regulatory network that controls black scale development in the mid-forewing band in *Heliconius* [15,16]. These experiments suggested *WntA* controlled black scale development differently on the wings of *H. erato* and *H. melpomene*. More precisely, the CRISPR/Cas9 *WntA* mutant phenotypes highlighted a more restricted area of black scales affected in the forewing of *H. melpomene* compared to *H. erato*. This result clearly demonstrated that although *H. erato* and *H. melpomene* exhibit strikingly similar mid-forewing band patterns, the genetic architecture underlying their resemblance may be more different than previously thought. These differences likely result from changes in the expression and interactions between other factors of the gene regulatory network that control the distribution of black scales. A direct consequence of this finding is that for a perfect pattern match to evolve between *H. erato* and *H. melpomene*, adaptive changes might need to occur at additional loci, apart from *WntA*. However, thus far, the evolutionary consequences of these differences in the gene regulatory network between *H. erato* and *H. melpomene* have not been extensively tested.

Here, we precisely determine similarity of the mid-forewing pattern between 14 distinct co-mimetic *H. erato* and *H. melpomene* populations. Using quantitative measurements, we first show that all co-mimetics exhibit consistent differences in their mid-forewing colour pattern shapes. Next, using published *WntA* CRISPR/Cas9 KO phenotypes, we tested if these differences could be explained by *WntA* function. Our data demonstrate the existence of species-specific developmental constraints that limit the ability of selection to produce perfectly identical phenotypes. The phenotypic manifestation of these developmental constraints are a direct consequence of selection on different genetic backgrounds that evolved during 12-14 MY of independent evolution. We conclude that selection has not been able to rewire identical gene regulatory networks in *H. erato* and *H. melpomene* but has found an alternative route to drive the evolution of similar phenotypes, albeit not completely perfect.

## 2. Materials

### (a) Sampling and landmark analysis

We obtained 8 to 14 images of each of 14 mimicking races of *H. erato* and *H. melpomene* (Table S1–2). Images were obtained through the authors’ collections and collections made publicly available by Cuthill *et al.* 2019 [22] and Jiggins *et al.* 2019 [23]. Individual genders were determined based on sexual dimorphism in the androconial region [24]. Landmarks were placed at 18 wing vein intersections on one forewing of each individual using *ImageJ* [25]. Landmarks 1, 6 and 10-18 were used in all analysis. Landmarks 2-5 and 7-9 were removed in a subset landmark analysis as they showed allometric shape differences between *H. erato* and *H. melpomene*. Landmarks were superimposed using Procrustes superimposition with the *procSym* function in the R package *Morpho* [26]. This superimposition transforms the raw landmark coordinates to a common centroid, scaling to unit centroid size, and rotating the shapes until the sum of squared distances between landmarks is minimized. The resulting Procrustes coordinates then describe shape differences between the samples. Tension maps represent the Euclidean distance between the average *H. erato* and *H. melpomene* Procrustes landmark arrangement and were created with a modified *tps_iso* and *tps_arr* function of the R package *Momocs* [27]. Landmark Principal Component Analysis (PCA) was performed with the *procSym* function in the R package *Morpho* [26] and ignoring size differences between wings (i.e. *sizeshape = FALSE*).

### (b) Color pattern analysis

Mid-forewing band patterns were extracted and aligned using the R package *patternize* [28]. Note that *WntA* defines black scale development in *Heliconius* [29], but for analysis purposes and because we were interested in variation of the mid-forewing band, we extracted and focused on the area of the forewing in which *WntA* is not expressed. Depending on the mid-forewing band phenotype, we specified RGB values for red, yellow and/or white with a color offset threshold (*colOffset*) chosen to fully extract the pattern. For *H. e. notabilis, H. m. plesseni* and *H. m. cythera*, we extracted and combined both red and white to represent the mid-forewing band shape. Background noise or damaged regions in the wing that were co-extracted with the color patterns were masked using the *setMask* function. Next, using *patternize* [28], a thin plate spline (tps) transformation was obtained from transforming landmarks to a common reference sample. This common reference sample included the landmarks of an arbitrarily chosen sample and was used as reference in all color pattern analysis. The tps transformation was then used to align and compare the extracted mid-forewing band shapes, size and position.

Differences in the mid-forewing band patterns were first compared by subtracting the average *H. erato* and *H. melpomene* mid-forewing band pattern of each population, obtained with the *sumRaster* function in *patternize* (i.e. absolute size difference). Next, the relative size of the mid-forewing band pattern was calculated as the proportion of the total wing area in which the pattern is expressed, using the *patArea* function (i.e. relative size difference). Principal Component Analysis (PCA) was performed on the binary representation of the aligned color pattern rasters obtained from each sample [28]. The PCA visualizes the main variations in color pattern boundaries among samples and groups and provides predictions of color pattern changes along the principal component (PC) axis. In the visualization of the predicted color pattern changes along the PC axis, positive values present a higher predicted expression of the pattern, whereas negative values present the absence of the pattern. Parts of the color patterns that are expressed in all samples have a predicted value of zero, as these pixels do not contribute variance for the PC analysis.

### (c) WntA CRISPR KO analysis

Five mutant butterflies of the Panamanian geographic races *Heliconius erato demophoon* and *Heliconius melpomene rosina* for which a frame shift mutation was generated at the gene *WntA* using CRISPR/Cas9 were obtained from Concha *et al.* (2019) [5]. All these mutants showed symmetric changes in wing patterns on both the left and right forewings and were thus likely full KO mutants [5]. Both left and right forewings were landmarked. Red was extracted from the *H. e. demophoon* mutants. As the *H. m. rosina* mutants often showed a yellow spot appearing in the proximal part of the mid-forewing band, both red and yellow were extracted for *H. m. rosina*. The color pattern expressed in the forewing of the mutants was extracted and aligned using the R package *patternize* [28]. The 90% quantile of the mutant pattern expression was obtained using the c*ontour* function of the R package *raster* [30] and superimposed on the wildtype wing pattern comparisons by aligning to the common reference sample.

## 3. Results

### (a) Controlling for allometric changes in wing shape and sex differences

Allometric differences in wing shape could potentially affect color pattern comparisons by overcompensating the pattern alignment compared to its relative position in the wing. Therefore, in our downstream color pattern analysis, we used two sets of landmarks: (1) one with all 18 vein intersection points and (2) and a second analysis excluding landmarks that caused allometric tension in the alignment (Figure 1). Principal Component Analysis (PCA) of the complete set of 18 landmarks placed at the intersection of every wing vein in the total set of 281 samples showed shape differences between *H. erato* and *H. melpomene* wings (Figure 1). We therefore used tension maps to better visualize these allometric wing shape changes and their effect on the alignment between the two species (Figure 1B). Allometric shape differences were most apparent at landmarks 2, 3, 7, 8 and 9 and affected mostly the alignment at the bottom proximal to medial area and the top medial to distal area of the wing (red areas in top part Figure 1B). This allometric effect was largely removed by occluding landmarks 2-5 and 7-9. In this subset landmark alignment, only small allometric tension areas remain in the alignment (green areas in bottom part Figure 1B). Comparing wing shape between males and females showed no apparent difference in sex in both *H. erato* and *H. melpomene* (Figure 1C).

**Figure 1.**
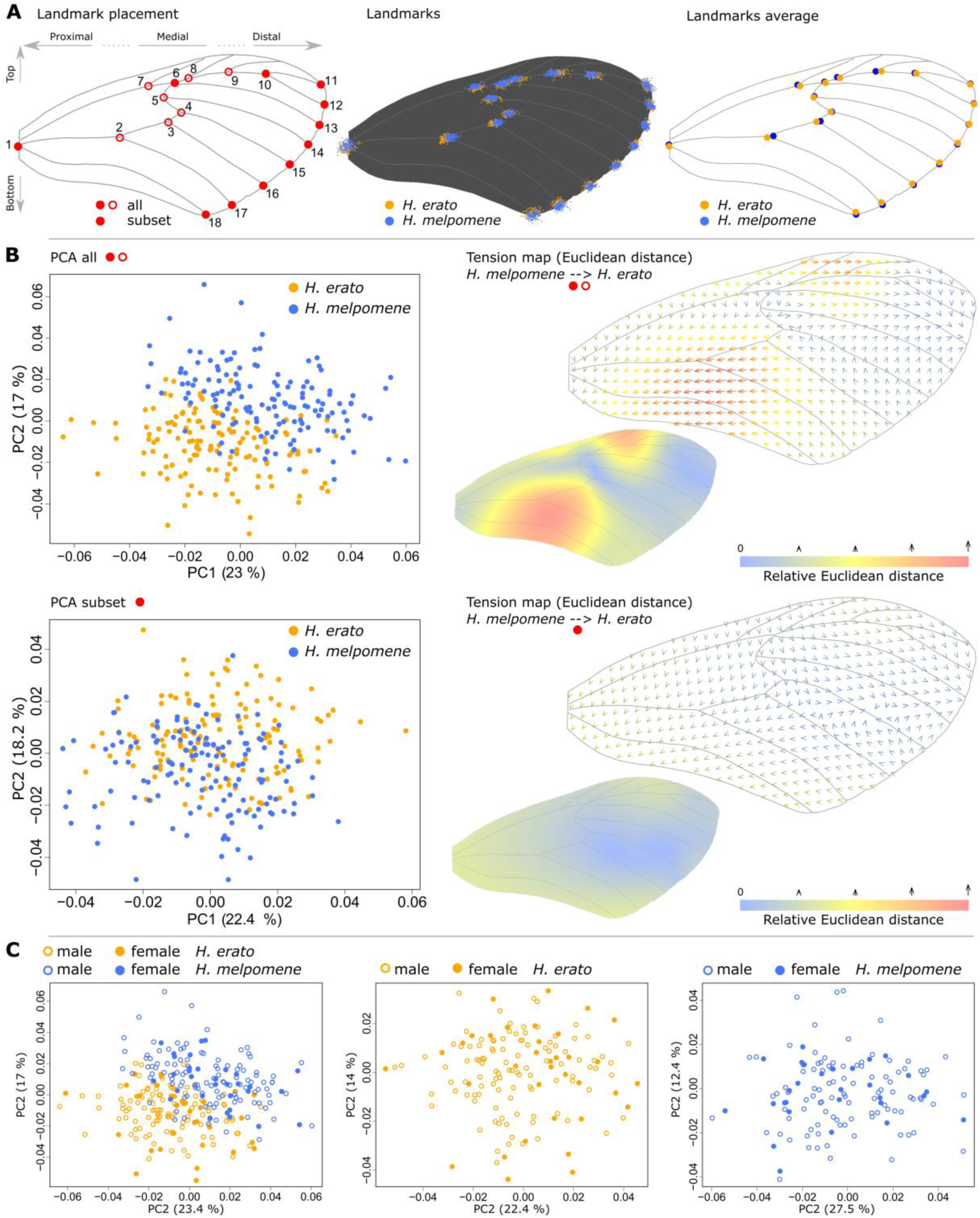
Landmark analysis shows allometric shape differences between *Heliconius erato* and *Heliconius melpomene* forewings but no differences between sexes. **(A)** Landmark placement of all 18 (open and closed circles) and subset (closed circles) (left), all landmarks placed in *H. erato* and *H. melpomene* samples (middle) and average landmark placement of *H. erato* and *H. melpomene* landmarks (right). **(B)** PCA of wing shape differences based on landmark placement (all 18 on top, subset in bottom) and tension in thin plate spline alignment. Colors and arrows in the tension map indicate allometric changes between *H. erato* and *H. melpomene*. Based on this analysis, we used the landmark subset which should not add any potential allometric alignment bias to the color pattern comparison. **(C)** Comparison of male (open circles) and female (closed circles) wing shape shows no differences in *H. erato* and *H. melpomene*.

### (b) Divergence and convergence in mid-forewing band pattern in H. erato and H. melpomene

In order to precisely quantify the similarity in mid-forewing band color pattern between *H. erato* and *H. melpomene* we sampled and analyzed a total of 281 images of 14 co-mimetic populations from Central and South America (Figure 2A, B). Geographic butterfly races cover a wide spectrum of mid-forewing band patterns, with unique or partially overlapping pattern elements among them. Interestingly, no populations that express the full potential mid-forewing band area exist in nature (Figure 2C). In the PCA of the mid-forewing band, the first main axis of variance (PC1) is dominated by the absence or presence of a broad red mid-forewing band, also typically called a ‘Postman’ phenotype (resembling the black and red uniforms worn by the Trinidad postal service; Figure 2C). The second main axis of variation (PC2) in the mid-forewing band shape is dominated by the presence of either a narrow median band or two spots as observed in the *H. e notabilis, H. e. microclea, H. m. plesseni* and *H. m. xenoclea* populations (Figure 2C).

**Figure 2.**
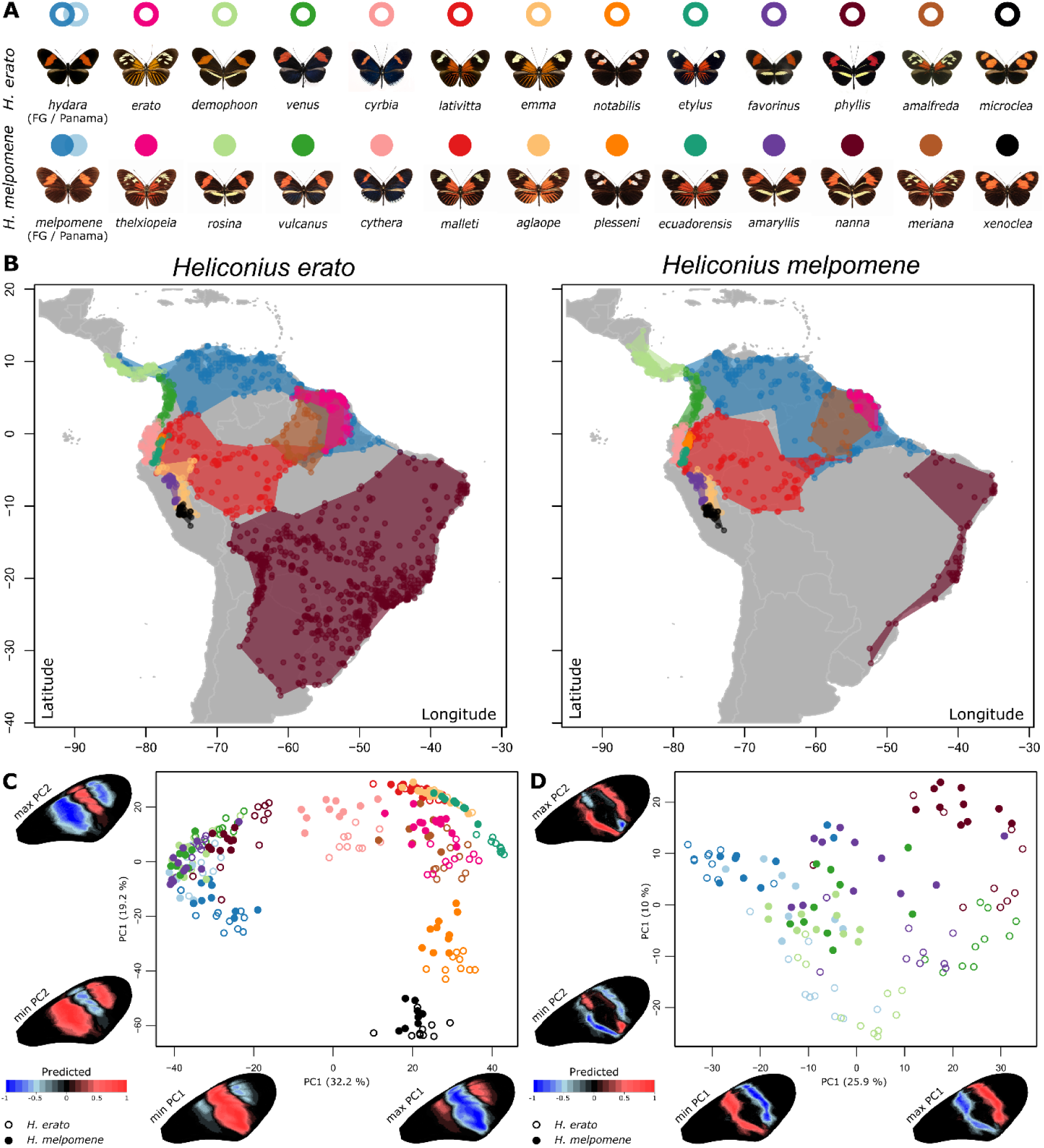
Mimicking *Heliconius erato* and *Heliconius melpomene* races, their distribution and PCA of mid-forewing band pattern. **(A)** Dorsal images of mimicking *H. erato* and *H. melpomene* races. **(B)** Distribution areas of the mimicking *H. erato* and *H. melpomene* races as obtained from Rosser *et al.* 2014 [48]. **(C)** PCA of mid-forewing band shape of the mimicking *H. erato* and *H. melpomene* races. **(D)** PCA of red banded ‘Postman’ races of mimicking *H. erato* and *H. melpomene* races. Wing heatmaps indicate minimum and maximum predicted patterns along each PC axis while considering the PC value of all other PC axis at zero. Positive values present a higher predicted expression of the pattern (red), whereas negative values present the absence of the pattern (blue).

As expected, co-mimicking races of *H. erato* and *H. melpomene* are found to have more similar mid-forewing band phenotypes than different populations of the same species. However, clear differences in clustering can be observed in the PCA between each of the co-mimicking pairs (Figure 2C, D), which reflect widespread mismatches across the mid-forewing band (Figure 3). These results thus indicate imperfection in the mimetic patterns between *H. erato* and *H. melpomene*, evolved to deter their predators. First, among co-mimicking races with a large medial red mid-forewing band (i.e. ‘Postman’ phenotypes), we observed an area in the distal bottom part of the mid-forewing band that consistently shows the absence of black scales in *H. melpomene* races compared to *H. erato* races (‘P’ in Figure 3). The consistency of this difference between *H. erato* and *H. melpomene* Postman races is also demonstrated by the second PC axis that describes mid-forewing band differences (Figure 2D). The second PC axis that describes mid-forewing band differences between *H. erato* and *H. melpomene* also demonstrates a general trend of expansion of both the proximal and distal area of the mid-forewing band in *H. melpomene* compared to *H. erato*. In more detailed comparisons among geographic Postman races, we generally observed an expansion of the proximal area of the mid-forewing band in *H. melpomene* compared to *H. erato* (Figure 3). However, this trend was reversed between the Postman races *H. e. hydara* and *H. m. melpomene* in French Guiana. In Colombia, the Postman races showed an expansion of the mid-forewing band in the distal area in *H. e. venus* compared to *H. m. vulcanus* (Figure 3). In both the latter cases, this variability in the pattern mismatch between the co-mimics can be largely ascribed by marked changes between the mid-forewing band of the *H. erato* Postman races *H. e. hydara* from Panama, *H. e. hydara* from French Guiana, and *H. e. venus*, with less pronounced phenotypic change among the *H. melpomene* Postman races *H. m. melpomene* from Panama, *H. m. melpomene* from French Guiana, and *H. m. vulcanus* (Figure 2D). Hence, these results highlight substantial divergence in wing color pattern between populations that are generally considered identical within *H. erato*.

**Figure 3.**
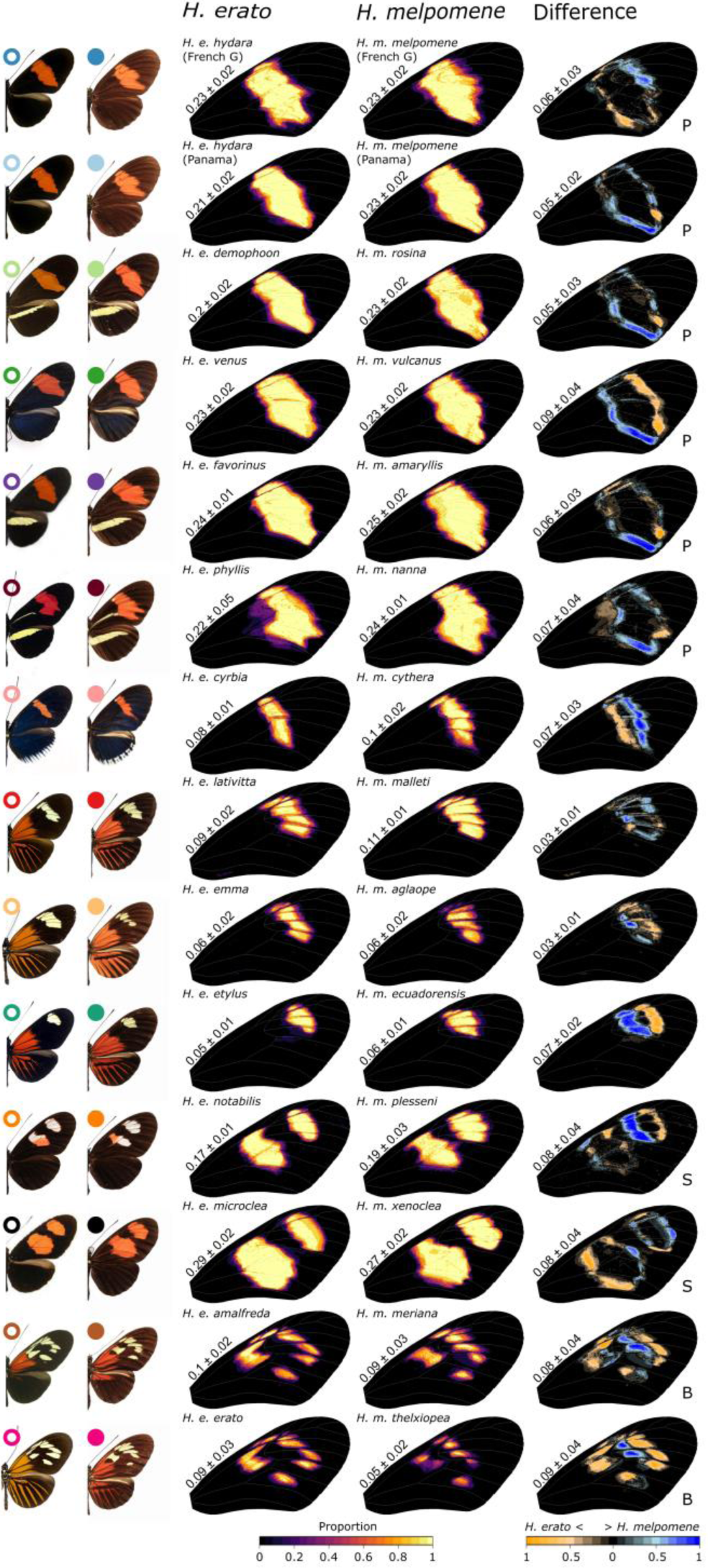
Quantification of mid-forewing band pattern in co-mimicking *Heliconius erato* and *Heliconius melpomene* races. Heatmaps demonstrate the consistency of mid-forewing band shape within races with light yellow indicating consistent expression and red and darker colors indicating less consistent expression among individuals. Differences in mid-forewing band shape between *H. erato* and *H. melpomene* races are shown on the right, with orange indicating higher expression of the trait in *H. erato* and blue indicating higher expression in *H. melpomene*. Values next to wings indicate the average proportion of the wing in which the trait is expressed or in which differences are found between *H. erato* and *H. melpomene* races. ‘P’ indicates phenotypes commonly referred to as the Postman races. ‘B’ indicates phenotypes commonly referred to as the Broken band races. ‘S’ indicates phenotypes commonly referred to as the Split band races. Color differences were quantified from aligned images using *patternize* [28] and results are robust to allometric changes in wing shape and sex differences (Figure S2, S3). Colored circles next to butterfly wing images correspond to distribution areas in Figure 2.

As observed in the Postman races, all other co-mimicking *Heliconius* races that we compared showed marked differences in their mid-forewing band (Figure 3). Comparison of the similarly patterned *H. e. cyrbia* with *H. m. cythera* from West Ecuador suggests that the position of the mid-forewing band is shifted proximally in *H. e. cyrbia* compared to *H. m. cythera*. In East Ecuador, the *H. m. malleti* samples investigated generally showed a larger mid-forewing band than the co-mimetic *H. e. lativitta* populations. Interestingly, differences in the comparison between the co-mimics *H. e. etylus* and *H. m. ecuadorensis*, which have a single distal spot, resembled differences in the distal spot between *H. e. notabilis* and *H. m. plesseni*, that have the so-called Split forewing band phenotype, which consists of two white/red spots in the mid-forewing band area (‘S’ in Figure 3). This finding suggests there may be similar developmental constraints in *H. e. etylus*/*H. e. notabilis* and *H. m. ecuadorensis*/*H. m. plesseni*. Finally, the so-called Broken mid-forewing band phenotypes *H. e. erato* and *H. e. amalfreda*, which consist of multiple yellow spots in the mid-forewing band area (‘B’ in Figure 3)., consistently differed from the *H. m. thelxipeia* and *H. m. meriana* populations by a proximal shift of the distal margin of the Broken mid-forewing band. This suggests a common genetic architecture of the Broken band phenotype within *H. erato* and *H. melpomene*, but potential developmental differences between them.

Interestingly, in all co-mimetic comparisons of *H. erato* and *H. melpomene*, the average absolute difference in the mid-forewing band pattern was larger than the average difference in the relative size (i.e. proportion of the wing in which the pattern is expressed) of the mid-forewing band pattern (Figure 4). Significant differences in the size of the mid-forewing band were only observed between the co-mimics *H. e. hydara* and *H. m. melpomene* from Panama (p = 0.013), *H. e. demophoon* and *H. m. rosina* (p = 0.013), *H. e. cyrbia* and *H. m. cythera* (p = 0.009) and *H. e. erato* and *H. m. thelxiopea* (p = 0.001). Despite the allometric shape changes observed between *H. erato* and *H. melpomene* (Figure 1), wing color pattern alignments including all 18 landmarks showed very similar PCA clustering for all populations and phenotypes compared to the subset landmark analysis. This shows that differences in mid-forewing band phenotype between *H. erato* and *H. melpomene* are largely independent from allometric shape differences in their wings (Figure S1, S2). Similarly, removing females from our dataset, which did not show differences in wing shape and size from males in our analyses, did not change the results (Figure S3).

**Figure 4.**
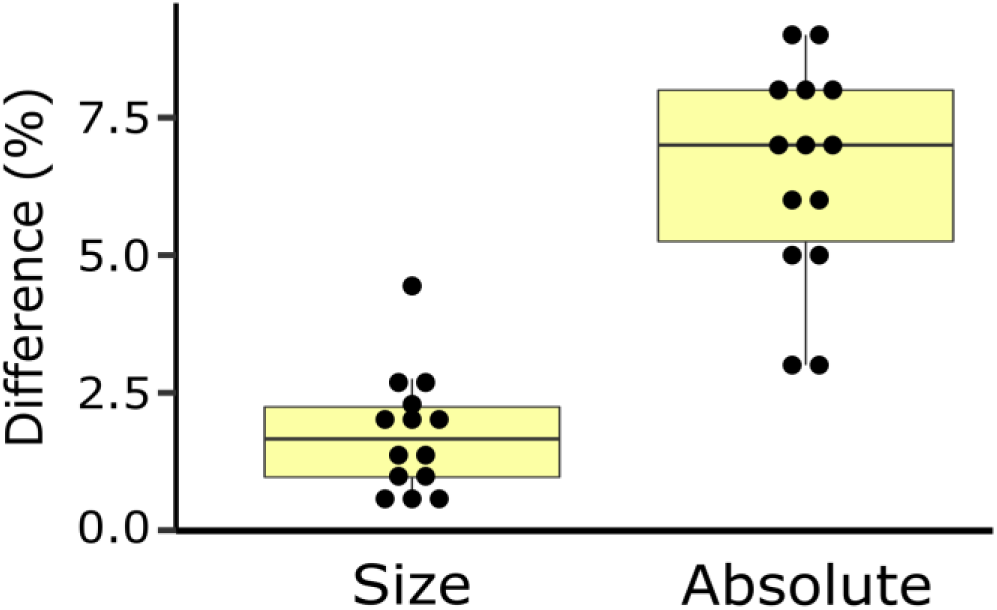
Difference in relative size and absolute difference of mid-forewing band pattern between *Heliconius erato* and *Heliconius melpomene*. Relative size indicates the size of the mid-forewing band pattern expressed as the percentage of the whole forewing area, without considering the position in the wing. Absolute difference indicates the mismatch in pattern when considering the exact position of the mid-forewing band within the wing. See Figure 3 for exact values.

### (c) Mismatch between co-mimics coincides with developmental WntA boundaries

As recently described by Concha *et al.* 2019 [5], *WntA* CRISPR/Cas9 KO’s of the Postman phenotype *H. e demophoon* showed a strong proximal expansion of the mid-forewing band pattern due to the development of red scales instead of black scales in this area of the wing (Figure 5A, left). In contrast, the *WntA* KO’s of the co-mimic *H. m. rosina* showed a less pronounced proximal expansion and an observable distal expansion of the mid-forewing band pattern (Figure 5A, right). Interestingly, in *H. m. rosina* a yellow spot sometimes appeared in the middle area of the proximal area of the mid-forewing band, suggesting different epistatic interactions in different parts of the wing in *H. melpomene* [5]. Aligning the expanded forewing area from the five *H. e demophoon* and five *H. m. rosina WntA* KO individuals confirms the marked different effect of *WntA* in the proximal area of the forewing (Figure 5B). An additional narrow difference is also observed at the distal margin of the mid-forewing band, with an expansion of the *WntA* affected area in *H. m. rosina* compared to *H. e. demophoon* (Figure 5B, top red arrows). At the bottom of this distal margin, this pattern is reversed and shows an area of the forewing where *WntA* affects scale coloration in *H. e. demophoon* but not in *H. m. rosina* (Figure 5B, bottom red arrows).

**Figure 5.**
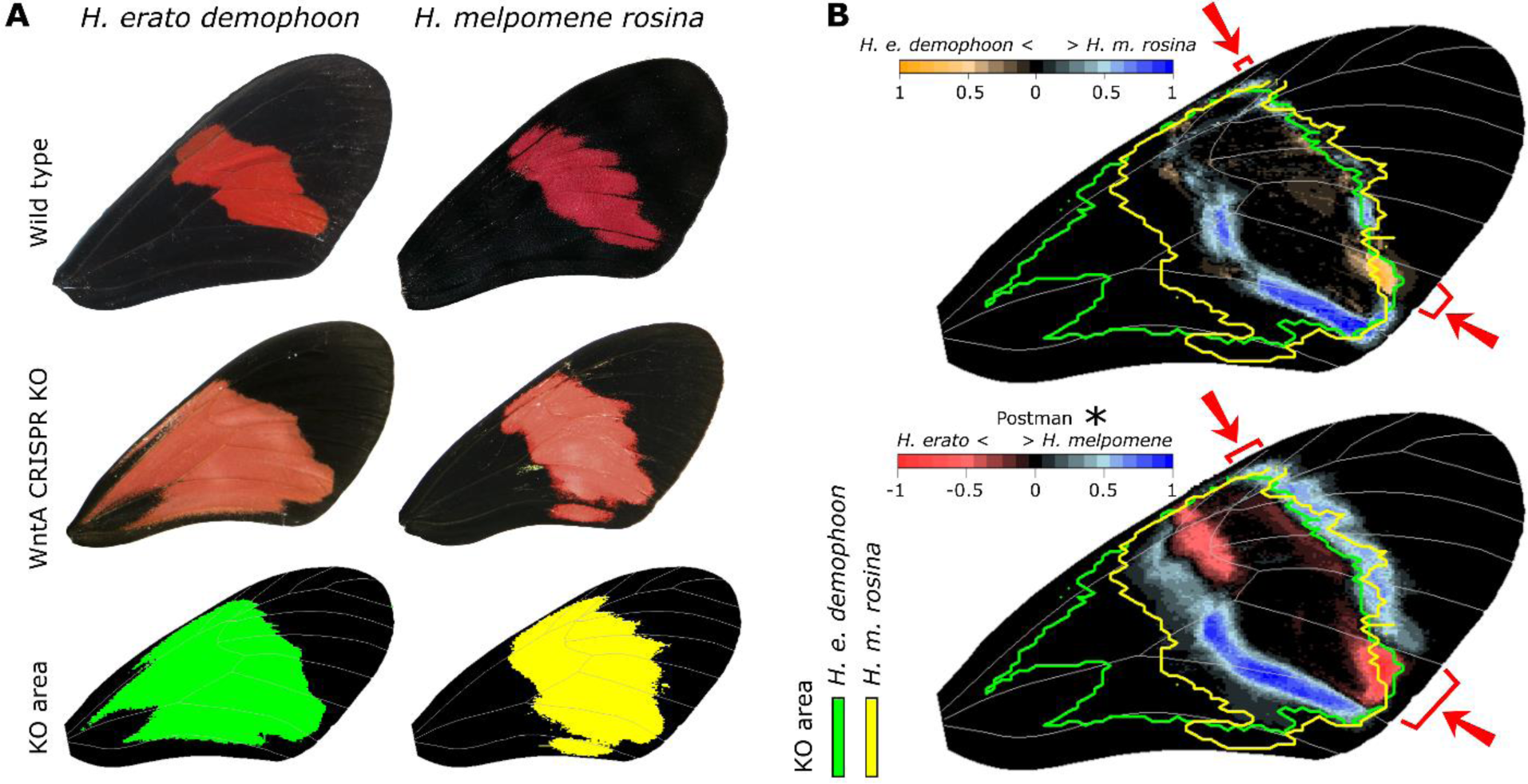
Mimetic differences between *Heliconius* butterflies demonstrate limits to adaptation imposed by developmental constraints. **(A)** Wild type, *WntA* CRISPR/Cas9 KO phenotypes (obtained from Concha *et al.* 2019 [5]) and 90% quantile of *WntA* CRISPR/Cas9 KO for *H. e. demophoon* and *H. m. rosina* (obtained from ten wings of five KO butterflies each). **(B)** Comparison of *WntA* CRISPR/Cas9 KO area to wild type difference between *H. e. demophoon* and *H. m. rosina* (top) and Postman race phenotype variation (bottom) (see Figure 2). Red arrows indicate overlap in mismatch between wild type mimicking populations and *WntA* KO patterning area.

The differences in the area affected by *WntA* in *H. e. demophoon* and *H. m. rosina* overlapped with a marked spot in which all *H. e. demophoon* wild type individuals differed from wild type *H. m. rosina* (Figure 5B, orange spot). A slight expansion of the *H. m. rosina* pattern compared to *H. e. demophoon* could also be observed within the distal area of differences between the *WntA* KO mutants. Mid-forewing band variation along the PC axis that mainly differentiated pattern variation between *H. erato* and *H. melpomene* Postman races recapitulated the distal spot difference (PC2 in Figure 1D). This result suggests that the distal forewing color pattern area that is generally expanded in *H. melpomene* Postman races perfectly matches the distal pattern boundary identified in the *H. e. demophoon WntA* mutants.

We also observed differences in the mid-forewing patterns between *H. erato* and *H. melpomene* mimics that overlap with the affected area seen in *WntA* mutants of both *H. e. demophoon* and *H. m. rosina* (Figure 3). These areas of mismatch that are within the boundaries of the *WntA* KO mutants may possibly indicate divergence in the regulatory architecture of the *WntA* gene. However, no formal genetic comparisons of the regulatory landscape near *WntA* have been performed so far between *H. erato* and *H. melpomene*. Finally, among the *H. erato* and *H. melpomene* mid-forewing band patterns investigated, none are found below the most proximal area in the forewing that is affected by *WntA* in *H. m. rosina*. The most proximal area of the forewing in which none-black scales are expressed is found in the Split band phenotypes *H. e. microclea* and *H. m. xenoclea*, at the vein intersection as marked by landmark 2 in Figure 1A. This observation suggests potential constraints on the forewing pattern diversity of these mimicking butterflies, constrained by the area of the wing that is controlled by *WntA* in *H. melpomene*.

## 4. Discussion

In *Heliconius* butterflies, convergence between co-mimicking populations has been broadly defined as nearly identical and has become one of the best visual examples of convergent evolution. In accordance with these phenotypic similarities, genetic work has highlighted that convergence is governed by a small and shared set of genes [6,15,17,21,31]. For example, the *WntA* gene has been repeatedly involved in wing color pattern convergence in the forewing of both *H. erato* and *H. melpomene* butterflies [15]. However, recent functional validation of *WntA* during wing development has provided new insights into the modality in which this gene controls patterning [5,21]. From these studies, *WntA* appears to affect a distinct wing color pattern domain in divergent co-mimetic butterflies [5]. This highlights the power of natural selection in driving convergent adaptive phenotypes despite a divergent genetic and developmental background, likely by independent changes in color pattern genes and possibly other uncharacterized genes [5]. However, to date, pattern similarity between co-mimetic *Heliconius* butterflies has been mostly described qualitatively. Therefore, we here compared differences observed between *WntA* KO’s of *H. erato* and *H. melpomene* with the quantitative variation found in the mid-forewing band of wild type co-mimetic butterflies to investigate to what extant divergence in the genetic and developmental background affects the evolution of identical phenotypes and if such differences correlate to *WntA* function.

### (a) Divergence in gene regulatory networks limits convergence

Convergence in wing color pattern between *H. erato* and *H. melpomene* has evolved through selection pressures imposed by birds that learn to associate the aposematic patterns with unpalatability [9]. From the bird predation, strong selection coefficients have been estimated for differences in forewing band patterns in *Heliconius* [for an overview see ref. 20]. Additionally, fine scale adjustments of the mid-forewing band in local butterfly communities have been identified in *Heliconius* [32]. Thus, it could be argued that there may be no limit in the ability to evolve perfect mimetic wing color patterns. However, our work clearly demonstrates that these *Heliconius* co-mimics are not as identical or perfect mimics as has been previous described. Mid-forewing band pattern differences were consistently found between all *H. erato* and *H. melpomene* populations investigated. Interestingly, our analyses show a strong correlation between these wild-type differences of *H. erato* and *H. melpomene* populations with differences in the *WntA* CRISPR/Cas9 KO phenotypes of *H. erato* and *H. melpomene* (Figure 4). This correspondence to the *WntA* KO boundaries demonstrates that the imperfections in mimicry are likely imposed by divergence in the network of other genes besides *WntA* that are involved in the development of the mid-forewing band. Alternatively, it is possible that differences in selection pressures, such as relaxed natural selection [8,33,34] or sexual selection [35], can explain imperfections in mimicry. However, our work highlights a strong correspondence of the differences in the developmental *WntA* boundaries between *H. erato* and *H. melpomene* that well explain the color pattern differences and thus demonstrate the importance of developmental constraints underlying the observed imperfect mimicry.

Divergence in the gene regulatory network that is involved in the development of the mid-forewing band may include both upstream factors that regulate spatial and/or temporal expression of *WntA* as well as genes that are downstream of *WntA* in the developmental pathway of melanic wing scales. A few of these diverged elements are likely loci or genes that have previously been implicated in wing color pattern variation in various *Heliconius* species. For example, the distal part of the mid-forewing band pattern that is expanded in *H. e. etylus*/*H. e. notabilis* compared to *H. m. ecuadorensis*/*H. m. plesseni* (Figure 3) matches the distal area of the wing that is described to be affected by an additional locus called *Ro*, described in *H. erato* over 30 years ago [16,36,37] and recently also identified to potentially affect the distal margin of the mid-forewing band in *H. melpomene* [38]. Next, differences in the mismatch between the Split band phenotypes *H. e. notabilis*/*H. m. plesseni* and *H. e. xenoclea*/*H. m. microclea* potentially result from epistatic interactions with the *optix* gene, affecting the size and shape of the mid-forewing band pattern. This has been previously suggested by looking at mid-forewing band patterns in hybrid butterflies that have the same *WntA* alleles but absence/presence of *optix* expression in the mid-forewing band [28]. Finally, contrasting results have been found regarding the number of loci that affect mid-forewing band shape in *H. erato* compared to *H. melpomene*. For example, in *H. erato* the genetic architecture of mid-forewing band pattern variation has been so far exclusively mapped to the so-called *Sd, St* and *Ly* loci that each affect a particular part of the mid-forewing band shape [36] and these loci have been demonstrated to include *cis*-regulatory elements of *WntA* [16]. In *H. melpomene*, on the other hand, regulatory variation at the so-called *N* locus, which likely includes the gene *cortex* or a closely linked gene, seems to also control mid-forewing band shape together with *WntA* [36]. Potentially, this latter locus provides a strong candidate that explains the restricted area in which *WntA* affects black wing scale development in *H. melpomene*.

### (b) Evidence of gene regulatory network divergence within species

*Heliconius* butterflies are also known for the incredibly diverse wing color pattern differences found in geographic races of the same species. Apart from the major effect loci involved in these color patterns, QTL studies of pattern variation in *H. erato* [39] and *H. melpomene* have demonstrated the existence of minor effect loci associated with quantitative changes in wing color pattern [38–40]. This larger set of genetic variants controlling quantitative variation is additional to the regulatory complexity that modulates the expression of the major color pattern genes [5,16,41]. However, these studies quantified pattern variation in crosses or hybrid zones between very distinct wing pattern phenotypes and did not directly compare geographically distinct populations that share similar wing color patterns. Here, we compared the shape, size and position of the mid-forewing band between populations of *H. erato* with a coloration that is generally described as the same general ‘Postman’ phenotype and found evidence for several quantitative differences (Figure 2D). Notably, the greatest mid-forewing band changes were observed between *H. erato* Postman races that also show the greatest imperfect mimicry with *H. melpomene* races (i.e. *H. e. hydara* versus *H. m. melpomene* from French Guiana). As *H. erato* is often suggested to be the more abundant co-mimic and, thus, the model which *H. melpomene* mimics [21,22], the evolution of better mimetic signals in this case may reflect a “lag” in the evolution of better mimicry imposed by intrinsic developmental limitations of the *H. melpomene* populations. This inference is further supported by the signal of recent adaptive evolution (i.e. selective sweep signal) across the regulatory regions in the first intronic region of *WntA* of *H. e. hydara* populations from French Guiana, but not *H. e. hydara* populations from Panama, or *H. m. melpomene* populations from French Guiana [20]. The potentially recent evolutionary change of the *H. e. hydara* mid-forewing band phenotype from French Guiana also crosses the developmental boundary identified in the *H. e. demophoon* CRISPR/Cas9 KO’s, which *s*uggests the gene regulatory network that underlies this similar wing phenotype may be diverging even within the *H. erato* lineage (Figure 3).

### (c) Mid-forewing patterns match well in size, but not in position

Some additional observations and questions arise from our work. For example, is it necessary for *H. erato* and *H. melpomene* to perfectly mimic each other or may small differences in pattern not impact the warning of potential predators? In our comparisons we observed that even though the position of forewing pattern elements may not be perfectly identical between co-mimics, the relative amount of black versus red or yellow is generally more similar than the absolute difference (Figure 4). This improved match of the size of the mid-forewing band seems to result from compensatory pattern changes in the proximal margin of the mid-forewing band. They thus indicate fine scale pattern adaptation in non-homologous regions of the wing to obtain a better match in the shape of the pattern even though they have a slightly shifted relative position in the wings of *H. erato* and *H. melpomene*. Such compensatory evolutionary changes to the mid-forewing band patterns could potentially include changes in the regulatory architecture of *WntA* that are less restricted by genetic and developmental constraints, compared to changes to other genes regulatory network.

The compensatory evolution to obtain a more similar area of the mid-forewing band despite its mismatch in position may also suggest imperfect discrimination in the visual range of their predators [42], or the relative importance of overall features of color contrast distribution rather than the exact position of pattern elements. A remarkable example of this are the co-mimetic Ecuadorian butterflies *H. e. notabilis* and *H. m. plesseni* which both have red and white in their proximal forewing element but have the relative positions of white versus red color inversed. Notably, these wing color patterns are the results of complex epistatic interactions between *WntA* and other genes such as the transcription factor *optix*, which controls white and red scale development [6,17].

## 5. Conclusion

The extent to which evolutionary changes are consequential for future adaptation has been most elegantly studied using microbes. In these experiments, multiple generations can be relatively easily traced while exposed to contrasting selection pressures and the consequences of their adaptations can be investigated when these selection pressures are reversed [1]. In malaria, for example, a single mutation of large effect can confer drug resistance but has been shown to also favor additional evolution of epistatic mutations [43]. Consequently, these changes have been demonstrated to create an epistatic ratchet against reverse evolution towards the ancestral phenotype, with important consequences for resistance management strategies [44]. Alternatively, in nematodes, cross-species conservation of gene expression during early life-stages has provided strong evidence for developmental constraints on the evolution of this stage within this phylum and animal evolution in general [45]. In non-experimental studies, the effect of genetic constraints on the direction of evolution has been suggested from correlations between genetic covariances within populations and the direction of morphological trait variation between species [46]. However, from such comparative studies it is challenging to infer the extent to which these genetic covariances limit adaptation or potential convergence. *In Heliconius*, constraints in the convergence of phenotypes is here identified as the result of divergence in the gene regulatory network that interacts with the gene *WntA* during black wing scale development. These constraints likely exist because the evolution of an improved pattern match between *H. erato* and *H. melpomene* would require modifications to the expression of additional genes in the gene regulatory network of the trait. These genes may not have the regulatory elements or architecture to easily be detached from potential maladaptive effects and strong developmental interactions with other genes [47].

## Data accessibility

All code to build the figures, landmarks and images are available through GitHub repository https://github.com/patternize-projects/Heliconius_forewing_band.

## Acknowledgements

We kindly thank Ananda Martins, Owen McMillan, Joe Hanly, Carolina Concha, Tim Thurman and Jennifer Cuthill for sharing images and Chris Jiggins, Ian Warran, Patricio Salazar and Gabriela Montejo-Kovacevich for the earthcape repository (https://heliconius.ecdb.io). We thank Frederik Nijhout for helpful comments on the manuscript.

## Supplementary information

**Table S 1.**
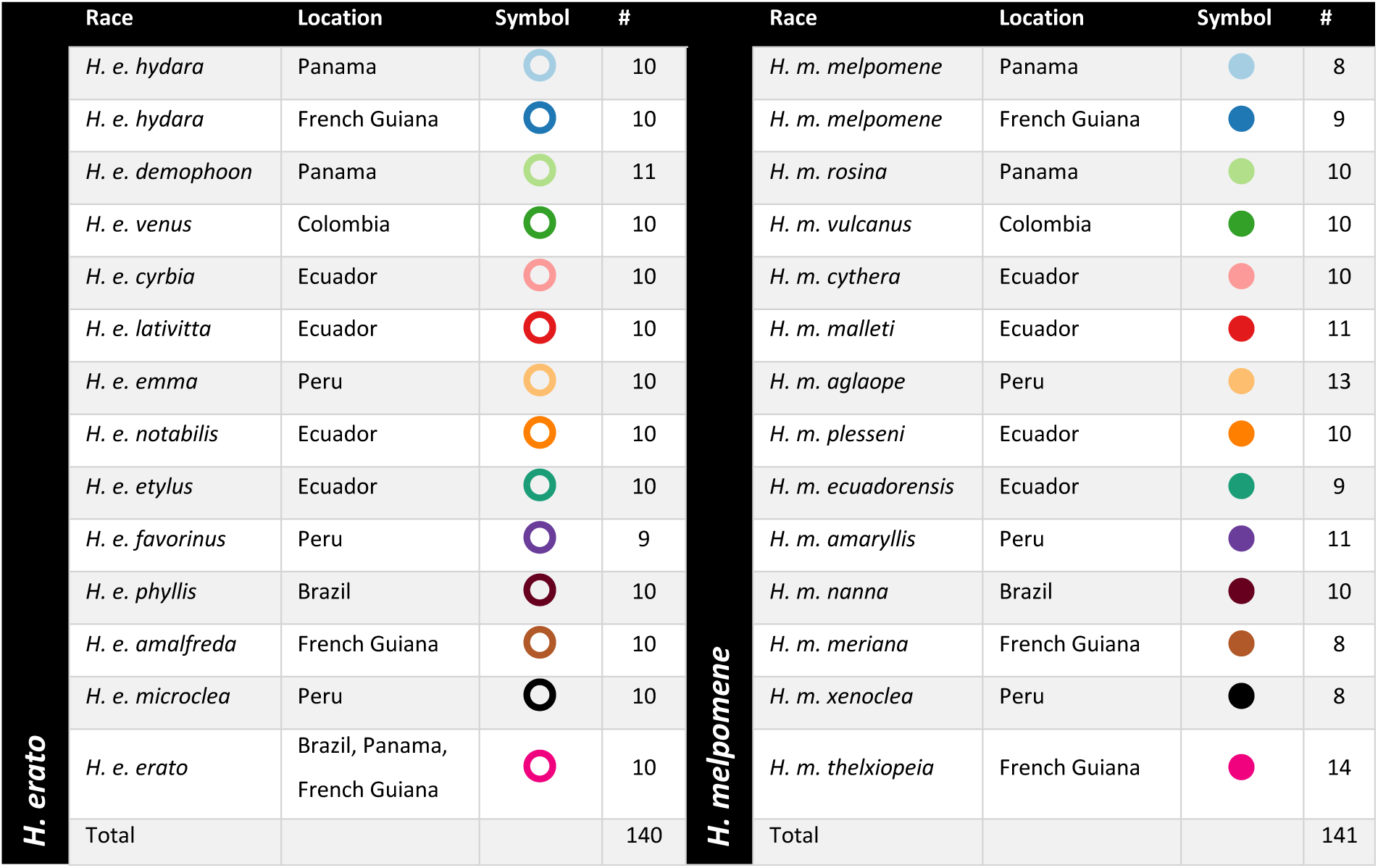
Summary of sampled images of each *H. erato* and *H. melpomene* race. Symbols are as in main Figure 2.

| *H. erato* | Race | Location | Symbol | # | *H. melpomene* | Race | Location | Symbol | # |
| --- | --- | --- | --- | --- | --- | --- | --- | --- | --- |
|  | *H. e. hydara* | Panama |  | 10 |  | *H. m. melpomene* | Panama |  | 8 |
|  | *H. e. hydara* | French Guiana |  | 10 |  | *H. m. melpomene* | French Guiana |  | 9 |
|  | *H. e. demophoon* | Panama |  | 11 |  | *H. m. rosina* | Panama |  | 10 |
|  | *H. e. venus* | Colombia |  | 10 |  | *H. m. vulcanus* | Colombia |  | 10 |
|  | *H. e. cyrbia* | Ecuador |  | 10 |  | *H. m. cythera* | Ecuador |  | 10 |
|  | *H. e. lativitta* | Ecuador |  | 10 |  | *H. m. malleti* | Ecuador |  | 11 |
|  | *H. e. emma* | Peru |  | 10 |  | *H. m. aglaope* | Peru |  | 13 |
|  | *H. e. notabilis* | Ecuador |  | 10 |  | *H. m. plesseni* | Ecuador |  | 10 |
|  | *H. e. etylus* | Ecuador |  | 10 |  | *H. m. ecuadorensis* | Ecuador |  | 9 |
|  | *H. e. favorinus* | Peru |  | 9 |  | *H. m. amaryllis* | Peru |  | 11 |
|  | *H. e. phyllis* | Brazil |  | 10 |  | *H. m. nanna* | Brazil |  | 10 |
|  | *H. e. amalfreda* | French Guiana |  | 10 |  | *H. m. meriana* | French Guiana |  | 8 |
|  | *H. e. microclea* | Peru |  | 10 |  | *H. m. xenoclea* | Peru |  | 8 |
|  | *H. e. erato* | Brazil, Panama,<br>French Guiana |  | 10 |  | *H. m. thelxiopeia* | French Guiana |  | 14 |
|  | Total |  |  | 140 |  | Total |  |  | 141 |

**Table S 2.** Sample ID, sex and localities. Images can be downloaded from https://github.com/patternize-projects/Heliconius_forewing_band.

| Species | Race | Image-ID | Sex | Latitude | Longitude | Location |
| --- | --- | --- | --- | --- | --- | --- |
| *H. erato* | *amalfreda* | BC2108 | m | 4.941100 | -55.182000 | Suriname |
| *H. erato* | *amalfreda* | BC2146 | m | 4.958700 | -55.179000 | Suriname |
| *H. erato* | *amalfreda* | BC2168 | f | 4.943490 | -55.186000 | Suriname |
| *H. erato* | *amalfreda* | BC2181 | m | 4.940650 | -55.190000 | Suriname |
| *H. erato* | *amalfreda* | BC2184 | f | 4.940650 | -55.190000 | Suriname |
| *H. erato* | *amalfreda* | BC2187 | m | 4.938030 | -55.194000 | Suriname |
| *H. erato* | *amalfreda* | BC2193 | f | 4.932730 | -55.201000 | Suriname |
| *H. erato* | *amalfreda* | BC2223 | f | 5.261830 | -55.622000 | Suriname |
| *H. erato* | *amalfreda* | BC2227 | m | 5.234000 | -55.775000 | Suriname |
| *H. erato* | *amalfreda* | BC2351 | f | 4.983940 | -55.156000 | Suriname |
| *H. erato* | *cyrbia* | CAM040321 | m | -0.052370 | -78.766000 | Ecuador |
| *H. erato* | *cyrbia* | CAM040354 | m | 0.207730 | -78.946910 | Ecuador |
| *H. erato* | *cyrbia* | CAM040355 | m | 0.207730 | -78.946910 | Ecuador |
| *H. erato* | *cyrbia* | CAM040356 | m | 0.207730 | -78.946910 | Ecuador |
| *H. erato* | *cyrbia* | CAM040357 | m | 0.207730 | -78.946910 | Ecuador |
| *H. erato* | *cyrbia* | CAM040361 | f | -0.052370 | -78.766000 | Ecuador |
| *H. erato* | *cyrbia* | CAM040362 | m | 0.140570 | -78.763030 | Ecuador |
| *H. erato* | *cyrbia* | CAM040363 | m | 0.140570 | -78.763030 | Ecuador |
| *H. erato* | *cyrbia* | CAM040364 | m | 0.140570 | -78.763030 | Ecuador |
| *H. erato* | *cyrbia* | CAM040365 | m | 0.140570 | -78.763030 | Ecuador |
| *H. erato* | *demophoon* | IMG_1960 | f | 9.166333 | -79.2079 | Panama |
| *H. erato* | *demophoon* | IMG_1972 | f | 9.166333 | -79.2079 | Panama |
| *H. erato* | *demophoon* | IMG_1974 | f | 9.166333 | -79.2079 | Panama |
| *H. erato* | *demophoon* | IMG_1978 | m | 9.2074 | -78.8265 | Panama |
| *H. erato* | *demophoon* | IMG_1980 | m | 9.2074 | -78.8265 | Panama |
| *H. erato* | *demophoon* | IMG_1982 | m | 9.2074 | -78.8265 | Panama |
| *H. erato* | *demophoon* | IMG_1987 | f | 9.2074 | -78.8265 | Panama |
| *H. erato* | *demophoon* | IMG_2049 | m | 9.2481 | -79.9478 | Panama |
| *H. erato* | *demophoon* | IMG_2125 | m | 9.11605 | -79.6984 | Panama |
| *H. erato* | *demophoon* | IMG_2135 | m | 9.11605 | -79.6984 | Panama |
| *H. erato* | *demophoon* | IMG_2141 | m | 9.11605 | -79.6984 | Panama |
| *H. erato* | *emma* | 10429077 | m | -1.583333 | -77.750000 | Ecuador |
| *H. erato* | *emma* | 10429078 | m | -1.583333 | -77.750000 | Ecuador |
| *H. erato* | *emma* | BC2563 | f | -5.29499 | -78.381 | Peru |
| *H. erato* | *emma* | BC2578 | m | -5.29499 | -78.381 | Peru |
| *H. erato* | *emma* | BC2579 | m | -5.29499 | -78.381 | Peru |
| *H. erato* | *emma* | BC2580 | m | -5.29499 | -78.381 | Peru |
| *H. erato* | *emma* | BC2611 | f | -5.38852 | -78.45114 | Peru |
| *H. erato* | *emma* | BC2612 | m | -5.38852 | -78.45114 | Peru |
| *H. erato* | *emma* | BC2620 | m | -5.37967 | -78.45478 | Peru |
| *H. erato* | *emma* | BC2624 | m | -5.44466 | -78.48361 | Peru |
| *H. erato* | *erato* | BC0147 | f | 4.622068 | -52.376361 | French Guiana |
| *H. erato* | *erato* | BC0148 | m | 4.556676 | -52.182277 | French Guiana |
| *H. erato* | *erato* | BC0149 | m | 4.607970 | -52.272347 | French Guiana |
| *H. erato* | *erato* | BC0154 | m | 4.563718 | -52.206697 | French Guiana |
| *H. erato* | *erato* | BC0163 | m | 4.568734 | -52.211198 | French Guiana |
| *H. erato* | *erato* | BC0198 | m | 4.570132 | -52.214164 | French Guiana |
| *H. erato* | *erato* | BC0200 | m | 4.564127 | -52.207655 | French Guiana |
| *H. erato* | *erato* | BC0327 | m | 4.638617 | -52.301596 | French Guiana |
| *H. erato* | *erato* | BC0340 | m | 4.570132 | -52.214164 | French Guiana |
| *H. erato* | *erato* | BC0351 | m | 4.548767 | -52.145497 | French Guiana |
| *H. erato* | *etylus* | 10429064 | m | 15.290278 | -89.146389 | Guatemala |
| *H. erato* | *etylus* | 10429062 | m | -1.583333 | -77.750000 | Ecuador |
| *H. erato* | *etylus* | BC3000 | f | -2.27322 | -78.19147 | Ecuador |
| *H. erato* | *etylus* | BC3001 | m | -2.27322 | -78.19147 | Ecuador |
| *H. erato* | *etylus* | BC3002 | m | -2.27322 | -78.19147 | Ecuador |
| *H. erato* | *etylus* | BC3003 | f | -2.27322 | -78.19147 | Ecuador |
| *H. erato* | *etylus* | BC3006 | f | -2.20024 | -78.15775 | Ecuador |
| *H. erato* | *etylus* | BC3007 | m | -2.20024 | -78.15775 | Ecuador |
| *H. erato* | *etylus* | BC3009 | f | -2.20024 | -78.15775 | Ecuador |
| *H. erato* | *etylus* | BC3010 | f | -2.20024 | -78.15775 | Ecuador |
| *H. erato* | *favorinus* | 10428999 | m | -9.300000 | -76.000000 | Peru |
| *H. erato* | *favorinus* | BC2634 | m | -6.4174 | -77.44329 | Peru |
| *H. erato* | *favorinus* | BC2635 | f | -6.4174 | -77.44329 | Peru |
| *H. erato* | *favorinus* | BC2636 | m | -6.4174 | -77.44329 | Peru |
| *H. erato* | *favorinus* | BC2637 | m | -6.4174 | -77.44329 | Peru |
| *H. erato* | *favorinus* | BC2638 | m | -6.4174 | -77.44329 | Peru |
| *H. erato* | *favorinus* | BC2640 | m | -6.4174 | -77.44329 | Peru |
| *H. erato* | *favorinus* | BC2643 | f | -6.41388 | -77.44505 | Peru |
| *H. erato* | *favorinus* | BC2646 | m | -6.41388 | -77.44505 | Peru |
| *H. erato* | *hydara* | BC0004 | m | 4.796800 | -52.324517 | French Guiana |
| *H. erato* | *hydara* | BC0049 | m | 4.796800 | -52.324517 | French Guiana |
| *H. erato* | *hydara* | BC0050 | m | 4.845894 | -52.348475 | French Guiana |
| *H. erato* | *hydara* | BC0061 | f | 4.549374 | -52.147880 | French Guiana |
| *H. erato* | *hydara* | BC0071 | m | 4.548767 | -52.145497 | French Guiana |
| *H. erato* | *hydara* | BC0076 | f | 4.796800 | -52.324517 | French Guiana |
| *H. erato* | *hydara* | BC0077 | m | 4.796800 | -52.324517 | French Guiana |
| *H. erato* | *hydara* | BC0079 | m | 4.796800 | -52.324517 | French Guiana |
| *H. erato* | *hydara* | BC0082 | m | 4.926844 | -52.390483 | French Guiana |
| *H. erato* | *hydara* | BC0125 | m | 4.616785 | -52.283083 | French Guiana |
| *H. erato* | *hydara* | IMG_1855 | m | 8.972917 | -78.5106 | Panama |
| *H. erato* | *hydara* | IMG_1861 | m | 8.972917 | -78.5106 | Panama |
| *H. erato* | *hydara* | IMG_1859 | m | 8.972917 | -78.5106 | Panama |
| *H. erato* | *hydara* | IMG_1863 | f | 8.950117 | -78.4496 | Panama |
| *H. erato* | *hydara* | IMG_1869 | m | 8.950117 | -78.4496 | Panama |
| *H. erato* | *hydara* | IMG_1871 | f | 8.972917 | -78.5106 | Panama |
| *H. erato* | *hydara* | IMG_1877 | m | 8.952533 | -78.5251 | Panama |
| *H. erato* | *hydara* | IMG_1885 | f | 8.972917 | -78.5106 | Panama |
| *H. erato* | *hydara* | IMG_1899 | m | 8.950117 | -78.4496 | Panama |
| *H. erato* | *hydara* | IMG_1993 | f | 9.2074 | -78.8265 | Panama |
| *H. erato* | *lativitta* | CAM016583 | m | -1.115567 | -77.778333 | Ecuador |
| *H. erato* | *lativitta* | CAM016586 | m | -1.115567 | -77.778333 | Ecuador |
| *H. erato* | *lativitta* | CAM016973 | f | -1.187817 | -77.831050 | Ecuador |
| *H. erato* | *lativitta* | CAM016999 | m | -1.187817 | -77.831050 | Ecuador |
| *H. erato* | *lativitta* | CAM017013 | f | -1.091567 | -77.719950 | Ecuador |
| *H. erato* | *lativitta* | CAM017027 | m | -1.091567 | -77.719950 | Ecuador |
| *H. erato* | *lativitta* | CAM017124 | m | -1.091567 | -77.719950 | Ecuador |
| *H. erato* | *lativitta* | CAM017170 | m | -1.061383 | -77.668433 | Ecuador |
| *H. erato* | *lativitta* | CAM017395 | f | -1.061383 | -77.668433 | Ecuador |
| *H. erato* | *lativitta* | CAM017405 | f | -1.098330 | -77.583890 | Ecuador |
| *H. erato* | *microclea* | 10428928 | m | -13.316667 | -71.600000 | Peru |
| *H. erato* | *microclea* | 10428929 | m | -9.883333 | -74.933333 | Peru |
| *H. erato* | *microclea* | 10428930 | m | -12.916667 | -71.400000 | Peru |
| *H. erato* | *microclea* | 10428931 | m | -11.066667 | -75.316667 | Peru |
| *H. erato* | *microclea* | 10428932 | m | -11.050000 | -75.316667 | Peru |
| *H. erato* | *microclea* | 10428934 | m | -10.883333 | -75.283333 | Peru |
| *H. erato* | *microclea* | 10428935 | m | -10.250000 | -74.750000 | Peru |
| *H. erato* | *microclea* | 10428936 | m | -10.883333 | -75.216667 | Peru |
| *H. erato* | *microclea* | 10428937 | m | -11.050000 | -75.316667 | Peru |
| *H. erato* | *microclea* | 10428941 | f | -11.066667 | -75.316667 | Peru |
| *H. erato* | *notabilis* | CAM016057 | m | -1.383017 | -77.944567 | Ecuador |
| *H. erato* | *notabilis* | CAM016058 | m | -1.383017 | -77.944567 | Ecuador |
| *H. erato* | *notabilis* | CAM016060 | f | -1.383017 | -77.944567 | Ecuador |
| *H. erato* | *notabilis* | CAM016067 | m | -1.362283 | -77.959433 | Ecuador |
| *H. erato* | *notabilis* | CAM016797 | f | -1.429250 | -77.986650 | Ecuador |
| *H. erato* | *notabilis* | CAM016894 | m | -1.437117 | -78.122883 | Ecuador |
| *H. erato* | *notabilis* | CAM016900 | m | -1.437117 | -78.122883 | Ecuador |
| *H. erato* | *notabilis* | CAM016915 | m | -1.459967 | -78.072800 | Ecuador |
| *H. erato* | *notabilis* | CAM017174 | m | -1.367483 | -78.016017 | Ecuador |
| *H. erato* | *notabilis* | CAM017177 | m | -1.367483 | -78.016017 | Ecuador |
| *H. erato* | *phyllis* | 10428488 | m | -16.833333 | -63.916667 | Bolivia |
| *H. erato* | *phyllis* | 10428490 | m | -17.250000 | -62.750000 | Bolivia |
| *H. erato* | *phyllis* | 10_MARA | m | -2.614010 | -44.283920 | Brasil |
| *H. erato* | *phyllis* | 112_BOQ2 | m | -2.805583 | -43.820889 | Brasil |
| *H. erato* | *phyllis* | 114_BOQ2 | m | -2.805583 | -43.820889 | Brasil |
| *H. erato* | *phyllis* | 35_SAG | m | -2.645028 | -44.140333 | Brasil |
| *H. erato* | *phyllis* | 49_SAG | f | -2.645028 | -44.140333 | Brasil |
| *H. erato* | *phyllis* | 52_SAG | m | -2.645028 | -44.140333 | Brasil |
| *H. erato* | *phyllis* | 6_MARA | m | -2.614010 | -44.283920 | Brasil |
| *H. erato* | *phyllis* | 85_IG | m | -2.756444 | -44.321639 | Brasil |
| *H. erato* | *venus* | CS000046 | m | -77.373333 | 3.958333 | Colombia |
| *H. erato* | *venus* | CS000047 | f | -77.373333 | 3.958333 | Colombia |
| *H. erato* | *venus* | CS000265 | f | -77.373333 | 3.958333 | Colombia |
| *H. erato* | *venus* | CS000275 | m | -77.373333 | 3.958333 | Colombia |
| *H. erato* | *venus* | CS000278 | m | -77.373333 | 3.958333 | Colombia |
| *H. erato* | *venus* | CS000284 | m | -77.373333 | 3.958333 | Colombia |
| *H. erato* | *venus* | CS000286 | m | -77.373333 | 3.958333 | Colombia |
| *H. erato* | *venus* | CS000288 | m | -77.373333 | 3.958333 | Colombia |
| *H. erato* | *venus* | CS003656 | m | -76.757222 | 3.500833 | Ecuador |
| *H. erato* | *venus* | CS003659 | m | -76.757222 | 3.500833 | Ecuador |
| *H. melpomene* | *agalope* | 10428229 | m | -4.273539 | -79.204614 | Ecuador |
| *H. melpomene* | *agalope* | 10428230 | m | -4.273539 | -79.204614 | Ecuador |
| *H. melpomene* | *agalope* | 10428231 | m | -1.583333 | -77.750000 | Ecuador |
| *H. melpomene* | *agalope* | 10428232 | m | -1.583333 | -77.750000 | Ecuador |
| *H. melpomene* | *agalope* | 10428233 | m | -1.583333 | -77.750000 | Ecuador |
| *H. melpomene* | *agalope* | 10428234 | m | -1.583333 | -77.750000 | Ecuador |
| *H. melpomene* | *agalope* | 10428235 | m | -1.583333 | -77.750000 | Ecuador |
| *H. melpomene* | *agalope* | 10428238 | f | -1.538392 | -78.067322 | Ecuador |
| *H. melpomene* | *agalope* | 10428240 | f | -1.066667 | -77.550000 | Ecuador |
| *H. melpomene* | *agalope* | 10428266 | m | -10.499805 | -75.652458 | Ecuador |
| *H. melpomene* | *agalope* | 10428267 | m | -13.433333 | -70.383333 | Ecuador |
| *H. melpomene* | *agalope* | CAM008689 | m | -6.289700 | -76.228900 | Peru |
| *H. melpomene* | *agalope* | CAM008702 | f | -6.239200 | -76.268700 | Peru |
| *H. melpomene* | *amaryllis* | 10428116 | m | -6.483333 | -76.366667 | Peru |
| *H. melpomene* | *amaryllis* | BC2639 | m | -6.417400 | -77.443290 | Peru |
| *H. melpomene* | *amaryllis* | MJ12.3137 | m | -6.454740 | -76.299440 | Peru |
| *H. melpomene* | *amaryllis* | MJ12.3371 | m | -6.452830 | -76.286215 | Peru |
| *H. melpomene* | *amaryllis* | MJ12.3392 | m | -6.452830 | -76.286215 | Peru |
| *H. melpomene* | *amaryllis* | MJ12.3393 | m | -6.452830 | -76.286215 | Peru |
| *H. melpomene* | *amaryllis* | MJ12.3396 | m | -6.452830 | -76.286215 | Peru |
| *H. melpomene* | *amaryllis* | MJ12.3414 | m | -6.454010 | -76.300230 | Peru |
| *H. melpomene* | *amaryllis* | MJ12.3416 | m | -6.453700 | -76.298070 | Peru |
| *H. melpomene* | *amaryllis* | MJ12.3417 | m | -6.453700 | -76.298070 | Peru |
| *H. melpomene* | *amaryllis* | MJ12.3442 | m | -6.454740 | -76.299440 | Peru |
| *H. melpomene* | *cythera* | 15N005 | f | 0.175330 | -78.907520 | Ecuador |
| *H. melpomene* | *cythera* | 15N006 | m | 0.193660 | -78.858700 | Ecuador |
| *H. melpomene* | *cythera* | 15N009 | m | 0.175330 | -78.907520 | Ecuador |
| *H. melpomene* | *cythera* | 15N020 | m | 0.184970 | -78.853020 | Ecuador |
| *H. melpomene* | *cythera* | 15N022 | f | 0.184970 | -78.853020 | Ecuador |
| *H. melpomene* | *cythera* | 15N023 | m | 0.184970 | -78.853020 | Ecuador |
| *H. melpomene* | *cythera* | CAM008510 | m | -0.648000 | -78.789500 | Ecuador |
| *H. melpomene* | *cythera* | CAM040383 | f | 0.212420 | -78.938550 | Ecuador |
| *H. melpomene* | *cythera* | CAM040459 | m | 0.149100 | -78.759070 | Ecuador |
| *H. melpomene* | *cythera* | CAM040474 | f | 0.153760 | -78.758970 | Ecuador |
| *H. melpomene* | *ecuadorensis* | 10428219 | m | -4.066667 | -78.966667 | Ecuador |
| *H. melpomene* | *ecuadorensis* | 10428220 | m | -1.583333 | -77.750000 | Ecuador |
| *H. melpomene* | *ecuadorensis* | CAM009112 | m | -4.043900 | -78.986100 | Ecuador |
| *H. melpomene* | *ecuadorensis* | CAM009113 | m | -4.043900 | -78.986100 | Ecuador |
| *H. melpomene* | *ecuadorensis* | CAM009114 | m | -4.043900 | -78.986100 | Ecuador |
| *H. melpomene* | *ecuadorensis* | CAM009115 | m | -4.043900 | -78.986100 | Ecuador |
| *H. melpomene* | *ecuadorensis* | CAM009119 | m | -4.043900 | -78.986100 | Ecuador |
| *H. melpomene* | *ecuadorensis* | CAM009120 | m | -4.043900 | -78.986100 | Ecuador |
| *H. melpomene* | *ecuadorensis* | CAM009141 | m | -2.251000 | -78.200000 | Ecuador |
| *H. melpomene* | *malleti* | 10428198 | m | -1.583300 | -77.750000 | Ecuador |
| *H. melpomene* | *malleti* | 10428199 | m | -1.583300 | -77.750000 | Ecuador |
| *H. melpomene* | *malleti* | CAM016144 | m | -1.416050 | -77.729017 | Ecuador |
| *H. melpomene* | *malleti* | CAM016267 | m | -1.250967 | -77.698850 | Ecuador |
| *H. melpomene* | *malleti* | CAM016609 | f | -1.115567 | -77.778333 | Ecuador |
| *H. melpomene* | *malleti* | CAM016610 | m | -1.115567 | -77.778333 | Ecuador |
| *H. melpomene* | *malleti* | CAM016611 | m | -1.115567 | -77.778333 | Ecuador |
| *H. melpomene* | *malleti* | CAM017049 | m | -1.168350 | -77.781117 | Ecuador |
| *H. melpomene* | *malleti* | CAM017064 | m | -1.168350 | -77.781117 | Ecuador |
| *H. melpomene* | *malleti* | CAM017070 | m | -1.168350 | -77.781117 | Ecuador |
| *H. melpomene* | *melpomene* | 10428369 | f | 5.500000 | -54.033300 | French Guiana |
| *H. melpomene* | *melpomene* | 10428371 | f | 5.400000 | -54.083300 | French Guiana |
| *H. melpomene* | *melpomene* | CAM000413 | m | 4.913800 | -52.359500 | French Guiana |
| *H. melpomene* | *melpomene* | CAM001422 | m | 4.963200 | -52.420000 | French Guiana |
| *H. melpomene* | *melpomene* | CAM008171 | f | 4.963200 | -52.420000 | French Guiana |
| *H. melpomene* | *melpomene* | CAM008215 | m | 4.789000 | -52.404000 | French Guiana |
| *H. melpomene* | *melpomene* | CAM008218 | m | 4.963200 | -52.420000 | French Guiana |
| *H. melpomene* | *melpomene* | CAM008863 | m | 7.756800 | -77.684100 | Panama |
| *H. melpomene* | *melpomene* | CAM008887 | m | 7.756800 | -77.684100 | Panama |
| *H. melpomene* | *melpomene* | CAM008888 | m | 7.756800 | -77.684100 | Panama |
| *H. melpomene* | *melpomene* | CAM008954 | m | 7.636200 | -78.189700 | Panama |
| *H. melpomene* | *melpomene* | CAM008955 | m | 7.636200 | -78.189700 | Panama |
| *H. melpomene* | *melpomene* | CAM008956 | m | 7.636200 | -78.189700 | Panama |
| *H. melpomene* | *melpomene* | CAM008957 | m | 7.636200 | -78.189700 | Panama |
| *H. melpomene* | *melpomene* | CAM008979 | m | 7.636200 | -78.189700 | Panama |
| *H. melpomene* | *melpomene* | CAM009316 | m | 4.963200 | -52.420000 | French Guiana |
| *H. melpomene* | *melpomene* | CAM009317 | m | 4.963200 | -52.420000 | French Guiana |
| *H. melpomene* | *meriana* | 13715 | m | 3.688300 | -54.082500 | Suriname |
| *H. melpomene* | *meriana* | melp_14-103 | m | 5.113892 | -54.990106 | Suriname |
| *H. melpomene* | *meriana* | melp_14-108 | m | 5.113892 | -54.990106 | Suriname |
| *H. melpomene* | *meriana* | melp_14-110 | m | 5.113892 | -54.990106 | Suriname |
| *H. melpomene* | *meriana* | melp_14-111 | m | 5.113892 | -54.990106 | Suriname |
| *H. melpomene* | *meriana* | melp_14-122 | m | 5.113892 | -54.990106 | Suriname |
| *H. melpomene* | *meriana* | melp_14-133 | f | 5.113892 | -54.990106 | Suriname |
| *H. melpomene* | *meriana* | melp_14-138 | f | 5.113892 | -54.990106 | Suriname |
| *H. melpomene* | *nanna* | LMCI_105-62 | m | -19.059167 | -40.139722 | Brasil |
| *H. melpomene* | *nanna* | LMCI_105-63 | m | -19.059167 | -40.139722 | Brasil |
| *H. melpomene* | *nanna* | LMCI_105-64 | f | -19.059167 | -40.139722 | Brasil |
| *H. melpomene* | *nanna* | LMCI_183-10 | m | -3.754444 | -40.910278 | Brasil |
| *H. melpomene* | *nanna* | LMCI_183-11 | m | -3.754444 | -40.910278 | Brasil |
| *H. melpomene* | *nanna* | LMCI_183-14 | m | -3.754444 | -40.910278 | Brasil |
| *H. melpomene* | *nanna* | LMCI_183-15 | m | -3.754444 | -40.910278 | Brasil |
| *H. melpomene* | *nanna* | LMCI_183-16 | m | -3.754444 | -40.910278 | Brasil |
| *H. melpomene* | *nanna* | LMCI_183-18 | f | -3.754444 | -40.910278 | Brasil |
| *H. melpomene* | *nanna* | LMCI_183-19 | m | -3.754444 | -40.910278 | Brasil |
| *H. melpomene* | *plesseni* | CAM016347 | f | -1.425867 | -78.018200 | Ecuador |
| *H. melpomene* | *plesseni* | CAM016349 | f | -1.425867 | -78.018200 | Ecuador |
| *H. melpomene* | *plesseni* | CAM016354 | m | -1.425867 | -78.018200 | Ecuador |
| *H. melpomene* | *plesseni* | CAM016355 | m | -1.425867 | -78.018200 | Ecuador |
| *H. melpomene* | *plesseni* | CAM016378 | m | -1.425867 | -78.018200 | Ecuador |
| *H. melpomene* | *plesseni* | CAM016810 | f | -1.409933 | -77.915350 | Ecuador |
| *H. melpomene* | *plesseni* | CAM017185 | m | -1.367483 | -78.016017 | Ecuador |
| *H. melpomene* | *plesseni* | CAM017186 | f | -1.367483 | -78.016017 | Ecuador |
| *H. melpomene* | *plesseni* | CAM017187 | f | -1.367483 | -78.016017 | Ecuador |
| *H. melpomene* | *plesseni* | CAM017614 | m | -1.422417 | -78.172933 | Ecuador |
| *H. melpomene* | *rosina* | CAM000903 | f | 9.129900 | -79.715800 | Panama |
| *H. melpomene* | *rosina* | CAM000947 | m | 9.129900 | -79.715800 | Panama |
| *H. melpomene* | *rosina* | CAM001015 | m | 9.116000 | -79.698000 | Panama |
| *H. melpomene* | *rosina* | CAM001027 | m | 9.076000 | -79.659000 | Panama |
| *H. melpomene* | *rosina* | CAM001067 | m | 9.010900 | -79.547700 | Panama |
| *H. melpomene* | *rosina* | CAM001137 | f | 9.122200 | -79.714900 | Panama |
| *H. melpomene* | *rosina* | CAM001391 | f | 9.122200 | -79.714900 | Panama |
| *H. melpomene* | *rosina* | CAM002901 | m | NA | NA | Panama |
| *H. melpomene* | *rosina* | CAM008052 | m | 8.981000 | -82.240000 | Panama |
| *H. melpomene* | *rosina* | CAM009554 | m | 8.975800 | -78.375300 | Panama |
| *H. melpomene* | *thelxiopea* | 10428329 | m | 5.250000 | -54.250000 | French Guiana |
| *H. melpomene* | *thelxiopea* | 10428331 | f | 5.400000 | -54.083333 | French Guiana |
| *H. melpomene* | *thelxiopea* | 10428332 | f | 5.400000 | -54.083333 | French Guiana |
| *H. melpomene* | *thelxiopea* | 10428333 | m | 5.400000 | -54.083333 | French Guiana |
| *H. melpomene* | *thelxiopea* | 10428334 | m | 5.400000 | -54.083333 | French Guiana |
| *H. melpomene* | *thelxiopea* | 10428335 | f | 5.400000 | -54.083333 | French Guiana |
| *H. melpomene* | *thelxiopea* | 10428336 | f | 5.400000 | -54.083333 | French Guiana |
| *H. melpomene* | *thelxiopea* | 10428337 | m | 5.400000 | -54.083333 | French Guiana |
| *H. melpomene* | *thelxiopea* | 10428338 | m | 5.250000 | -54.250000 | French Guiana |
| *H. melpomene* | *thelxiopea* | 10428339 | m | 5.500000 | -54.033333 | French Guiana |
| *H. melpomene* | *thelxiopea* | 10428340 | f | 5.400000 | -54.083333 | French Guiana |
| *H. melpomene* | *thelxiopea* | 10428341 | f | 5.250000 | -54.250000 | French Guiana |
| *H. melpomene* | *thelxiopea* | 10428342 | f | 5.516667 | -53.966667 | French Guiana |
| *H. melpomene* | *thelxiopea* | 10428343 | m | 5.500000 | -54.033333 | French Guiana |
| *H. melpomene* | *vulcanus* | CAM000058 | m | 3.777900 | -76.130000 | Colombia |
| *H. melpomene* | *vulcanus* | CAM000059 | m | 3.782500 | -76.721100 | Colombia |
| *H. melpomene* | *vulcanus* | CAM000060 | m | 3.782500 | -76.721100 | Colombia |
| *H. melpomene* | *vulcanus* | CAM000061 | m | 3.782500 | -76.721100 | Colombia |
| *H. melpomene* | *vulcanus* | CAM000062 | m | 3.782500 | -76.721100 | Colombia |
| *H. melpomene* | *vulcanus* | CAM000063 | m | 3.782500 | -76.721100 | Colombia |
| *H. melpomene* | *vulcanus* | CAM000064 | m | 3.782500 | -76.721100 | Colombia |
| *H. melpomene* | *vulcanus* | CAM000129 | m | 3.895800 | -76.621400 | Colombia |
| *H. melpomene* | *vulcanus* | CAM000132 | m | 3.895800 | -76.621400 | Colombia |
| *H. melpomene* | *vulcanus* | CAM000134 | m | 3.895800 | -76.621400 | Colombia |
| *H. melpomene* | *xenoclea* | MJ12.3605 | m | -11.174530 | -75.403470 | Peru |
| *H. melpomene* | *xenoclea* | MJ12.3606 | m | -11.174530 | -75.403470 | Peru |
| *H. melpomene* | *xenoclea* | MJ12.3608 | m | -11.174530 | -75.403470 | Peru |
| *H. melpomene* | *xenoclea* | MJ12.3636 | m | -11.033770 | -75.409130 | Peru |
| *H. melpomene* | *xenoclea* | MJ12.3638 | m | -11.036420 | -75.407990 | Peru |
| *H. melpomene* | *xenoclea* | MJ12.3647 | m | -11.044580 | -75.413270 | Peru |
| *H. melpomene* | *xenoclea* | MJ12.3648 | m | -11.033770 | -75.409130 | Peru |
| *H. melpomene* | *xenoclea* | MJ12.3651 | m | -11.033770 | -75.409130 | Peru |
| *H. melpomene* | *xenoclea* | MJ12.3653 | m | -11.033770 | -75.409130 | Peru |

**Figure S1.**
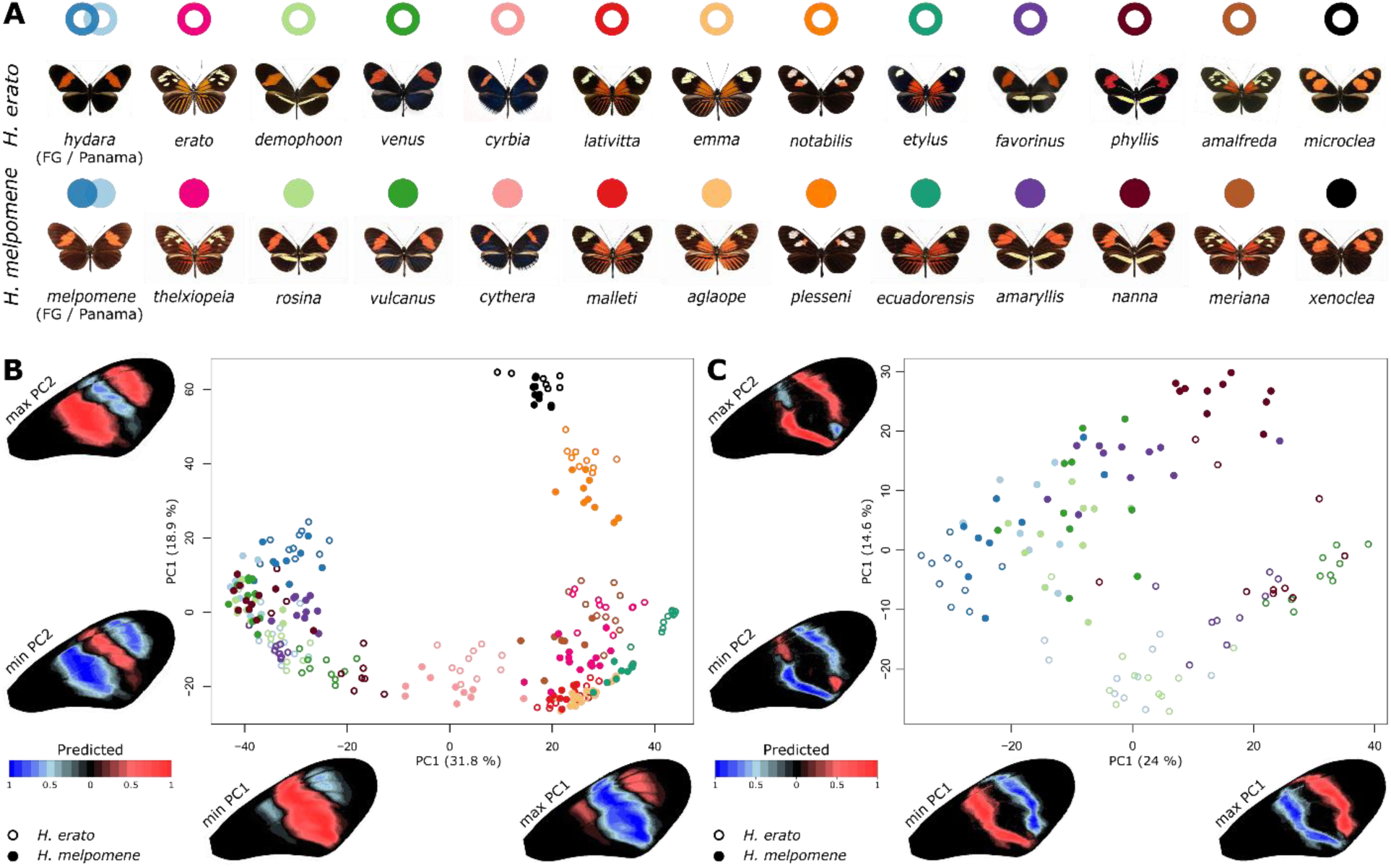
Principal Component Analysis (PCA) of mid-forewing band shape in *H. erato* and *H. melpomene* using all 18 landmarks. **(A)** Dorsal images of mimicking *H. erato* and *H. melpomene* races. **(B)** PCA of mid-forewing band shape of the mimicking *H. erato* and *H. melpomene* races. **(C)** PCA of red banded ‘Postman’ races of mimicking *H. erato* and *H. melpomene* races. Wing heatmaps indicate minimum (blue) and maximum (red) predicted pattern along each PC axis.

**Figure S2.**
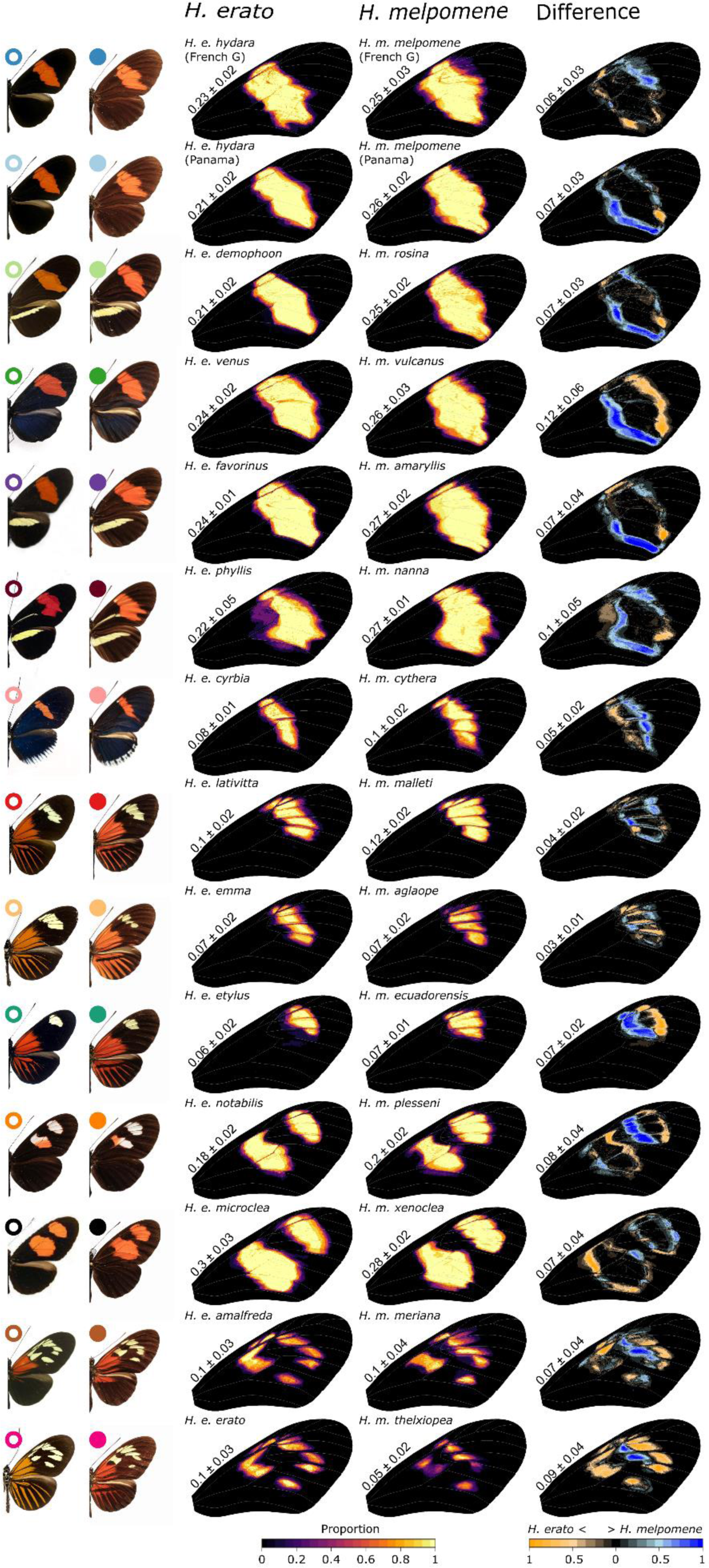
Quantification of mid-forewing band pattern in co-mimicking *H. erato* and *H. melpomene* races using all 18 landmarks. Heatmaps demonstrate the consistency of mid-forewing band shape within races with light yellow indicating consistent expression and red and darker colors indicating less consistent expression among individuals. Differences in mid-forewing band shape between *H. erato* and *H. melpomene* races are shown on the right, with orange indicating higher expression of the trait in *H. erato* and blue indicating higher expression in *H. melpomene*. Values next to wings indicate the average proportion of the wing in which the trait is expressed or in which differences are found between *H. erato* and *H. melpomene* races.

**Figure S3.**
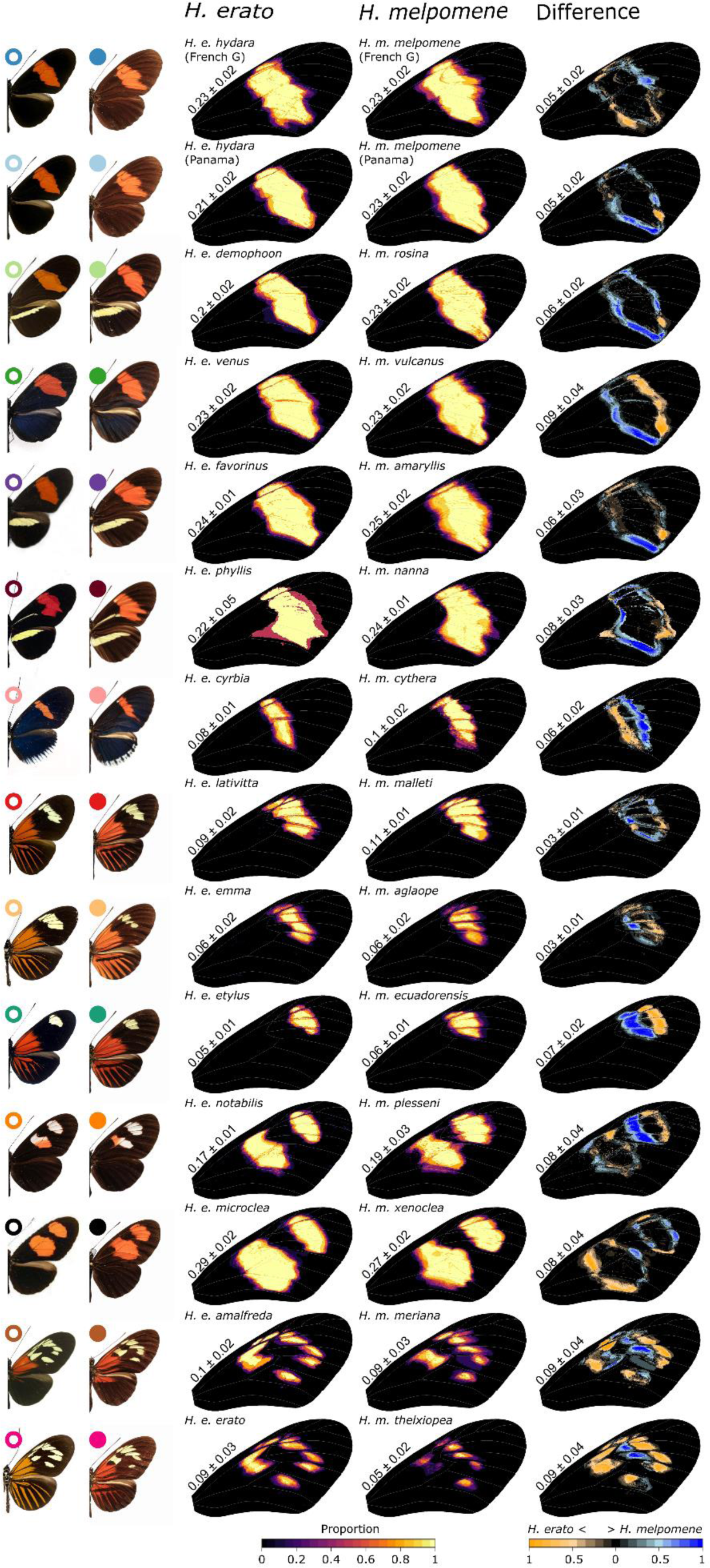
Quantification of forewing band pattern in co-mimicking *H. erato* and *H. melpomene* races using subset landmarks and only male samples. Heatmaps demonstrate the consistency of forewing band shape within races with light yellow indicating consistent expression and red and darker colors indicating less consistent expression among individuals. Differences in forewing band shape between *H. erato* and *H. melpomene* races are shown on the right, with orange indicating higher expression of the trait in *H. erato* and blue indicating higher expression in *H. melpomene*. Values next to wings indicate the average proportion of the wing in which the trait is expressed or in which differences are found between *H. erato* and *H. melpomene* races.

